# Dissociation of Medial Frontal β-Bursts and Executive Control

**DOI:** 10.1101/2020.08.06.240606

**Authors:** Steven P. Errington, Geoffrey F. Woodman, Jeffrey D. Schall

## Abstract

The neural mechanisms of executive and motor control concern both basic researchers and clinicians. In human studies, preparation and cancellation of movements are accompanied by changes in the β-frequency band (15–29 Hz) of EEG. Previous studies with human participants performing stop signal (countermanding) tasks have described reduced frequency of transient β-bursts over sensorimotor cortical areas before movement initiation and increased β-bursting over medial frontal areas with movement cancellation. This modulation has been interpreted as contributing to the trial-by-trial control of behavior. We performed identical analyses of EEG recorded over the frontal lobe of macaque monkeys performing a saccade countermanding task. Whilst, we replicate the occurrence and modulation of β-bursts associated with initiation and cancellation of saccades, we found that β-bursts occur too infrequently to account for the observed stopping behavior. We also found β-bursts were more common after errors, but their incidence was unrelated to response time adaptation. These results demonstrate the homology of this EEG signature between humans and macaques but raise questions about the current interpretation of β-band functional significance.

**SIGNIFICANCE STATEMENT:** The finding of increased β-bursting over medial frontal cortex with movement cancellation in humans is difficult to reconcile with the finding of modulation too late to contribute to movement cancellation in medial frontal cortex of macaque monkeys. To obtain comparable measurement scales, we recorded EEG over medial frontal cortex of macaques performing a stop signal (countermanding) task. We replicated the occurrence and modulation of β-bursts associated with the cancellation of movements, but we found that β-bursts occur too infrequently to account for observed stopping behavior. Unfortunately, this finding raises doubts whether β-bursts can be a causal mechanism of response inhibition, which impacts future applications in devices such as brain-machine interfaces.

## INTRODUCTION

Response inhibition and performance monitoring are two executive control functions supporting goal-directed behavior. The countermanding (stop-signal) task has provided numerous insights into these functions (Verbruggen and Logan, 2008; Verbruggen et al., 2019). Most recently, researchers recording human EEG during the stop signal task have described a higher incidence of β-bursts over medial frontal cortex during successful inhibition arising early enough to contribute to stopping the movement (Jana et al., 2020; Wessel, 2020). This result is difficult to reconcile with single-unit recordings from medial frontal cortex of macaques, which described neural signals modulating only after inhibition was achieved (Stuphorn et al., 2010) and most commonly after errors of response inhibition (Stuphorn et al., 2000; Sajad et al., 2019).

When applied to electrophysiological (De Jong et al., 1995; Kok et al., 2004; Stahl and Gibbons, 2007; Godlove et al., 2011b; Reinhart et al., 2012; Swann et al., 2012; Wessel and Aron, 2015) and neurophysiological (Hanes et al., 1998; Pare and Hanes, 2003; Brown et al., 2008; Stuphorn et al., 2010; Schmidt et al., 2013; Brockett et al., 2020) measurements, the model of a race between GO and STOP processes (Logan and Cowan, 1984) offers clear criteria to attribute neural activity to movement initiation and inhibition. To contribute to reactive response inhibition, a neural signal must appear before the STOP process finishes, and that signal must scale in frequency of occurrence proportional to the probability of canceling a response after a stop signal is presented. The spiking rate of movement-related neurons in cortical (Hanes and Schall, 1995; Murthy et al., 2009; Mirabella et al., 2011) and subcortical (Pare and Hanes, 2003; Schmidt et al., 2013) motor circuits satisfy these criteria, whilst the spiking rate of neurons in medial frontal areas do not (Scangos and Stuphorn, 2010; Stuphorn et al., 2010). Performance of the countermanding task typically includes failures to cancel the response on about half of all stop signal trials. This high failure rate also affords measurement and analysis of neural signals of error processing, which can interact with proactive control.

Analyzing EEG over medial frontal cortex of macaque monkeys performing a saccade stop signal task, we observed β-bursts as infrequently as found in previous human studies. We also observed slightly elevated proportions before the STOP process finished when responses were canceled. However, the probability of β-bursts lagged far behind the probability of canceling a response at each stop signal time. Indeed at least 80% of responses were canceled despite no β-burst occurring. Unexpectedly, we also observed pronounced increase in β-burst frequency after errors of inhibition, but the proportion of these β-bursts was unrelated to adaptation in response times (RT). We conclude that β-bursts may serve as a rough index of executive control processes, but they lack any causal efficacy and thus much theoretical pr practical utility.

## MATERIALS AND METHODS

### Experimental Model and Subject Details

All procedures were in accordance with the National Institutes of Health Guidelines, the American Association for Laboratory Animal Care *Guide for the Care and Use of Laboratory Animals* and approved by the Vanderbilt Institutional Animal Care and Use Committee in accordance with the United States Department of Agriculture and Public Health Service policies. Data was collected from one male bonnet macaque (Eu, *Macaca Radiata*, 8.8 kg) and one female rhesus macaque (X, *M. Mulatta*, 6.0 kg) performing a saccade countermanding task (Hanes and Schall, 1995; Godlove et al., 2014). Both animals were on a 12-hour light-dark cycle and all experimental procedures were conducted in the daytime. Each monkey received nutrient-rich, primate-specific food pellets twice a day. Fresh produce and other forms of environmental enrichment were given at least five times a week.

### Surgical Procedures

Surgical details have been described previously (Godlove et al., 2011a). Briefly, magnetic resonance images (MRIs) were acquired with a Philips Intera Achieva 3T scanner using SENSE Flex-S surface coils placed above or below the animal’s head. T1-weighted gradient-echo structural images were obtained with a 3D turbo field echo anatomical sequence (TR = 8.729 ms; 130 slices, 0.70 mm thickness). These images were used to ensure Cilux recording chambers were placed in the correct area (Crist Instruments). Chambers were implanted normal to the cortex (Monkey Eu: 17°; Monkey X: 9°; relative to stereotaxic vertical) centered on midline: 30 mm (Monkey Eu) and 28 mm (Monkey X) anterior to the interaural line.

### Data Collection Protocol

An identical daily recording protocol across monkeys and sessions was carried out. In each session, the monkey sat in an enclosed primate chair with their head restrained 45 cm from a CRT monitor (Dell P1130, background luminance of 0.10 cd/m^2^). The monitor had a refresh rate of 70 Hz, and the screen subtended 46 deg × 36 deg of the visual angle. Eye position data was collected at 1 kHz using an EyeLink 1000 infrared eye-tracking system (SR Research, Kanata, Ontario, Canada). All data were streamed to a single data acquisition system (MAP, Plexon, Dallas, TX). Time stamps of trial events were recorded at 500 Hz.

### Macaque Electroencephalography

The EEG was recorded from the cranial surface with an electrode located over medial frontal cortex. The electrode implants were constructed from Teflon-coated braided stainless-steel wire and solid-gold terminals. Implanted wires were cut to 8.5 cm, the wire ends exposed, and gold Amphenol pins were crimped to both ends. One end of the wires was inserted into a plastic connector, whereas the gold pin on the other end was ground down until 1 mm of the pin remained. During aseptic surgery, a ~1 mm hole was drilled into the surface of the skull (3–5 mm thick), allowing the terminal end of the electrode to be tightly inserted. The inserted gold pin was then covered with a small amount of acrylic cement. After the EEG electrode was implanted, the plastic connector was attached to exposed acrylic to allow access to the channels. Leads that were not embedded in the acrylic were covered by skin that was sutured back over the skull. This allowed for the EEG electrode to be minimally invasive once implanted. Unlike recordings from skull screws that extend to the dura mater through the skull, recordings from these electrodes approximate those used in human electrophysiological studies because the signals must propagate through the layers of brain, dura, and skull. Electrodes were referenced to linked ears using ear-clip electrodes (Electro-Cap International). The EEG from each electrode was amplified with a high-input impedance head stage (Plexon) and bandpass filtered between 0.7 and 170 Hz. All data were streamed to a data acquisition system (MAP, Plexon, Dallas, TX).

### Saccade Stop-Signal (Countermanding) Task

The saccade stop-signal task utilized in this study has been widely used previously (Hanes and Schall, 1995; Hanes and Carpenter, 1999; Cabel et al., 2000; Colonius et al., 2001; Kornylo et al., 2003; Morein-Zamir and Kingstone, 2006; Walton and Gandhi, 2006; Thakkar et al., 2011; Thakkar et al., 2015; Godlove and Schall, 2016; Wattiez et al., 2016; Verbruggen et al., 2019). Briefly, trials were initiated when monkeys fixated at a central point. Following a variable time period, the center of the fixation point was removed leaving an outline. At this point, a peripheral target was presented simultaneously on either the left or right hand of the screen.

In this study, one target location was associated with a larger magnitude of fluid reward. The lower magnitude reward ranged from 0 to 50% of the higher magnitude reward amount. This incidence was adjusted to encourage the monkey to continue responding to both targets. The stimulus-response mapping of location-to-high reward changed across blocks of trials. Block length was adjusted to maintain performance at both targets, with the number of trials in each block determined by the number of correct trials performed. In most sessions, the block length was set at 10 to 30 correct trials. Erroneous responses led to repetitions of a target location, ensuring that monkeys did not neglect low-reward targets in favor of high-reward targets – a phenomenon demonstrated in previous implementations of asymmetrically rewarded tasks (Kawagoe et al., 1998).

On most of the trials, the monkey was required to make an eye movement to this target (no-stop trials). However, on a proportion of trials the center of the fixation point was re-illuminated (stop-signal trials); this stop signal appeared at a variable time after the target had appeared (stop-signal delay; SSDs). An initial set of SSDs, separated by either 40 or 60 ms, were selected for each recording session. The delay was then manipulated through an adaptive staircasing procedure in which stopping difficulty was based on performance. When a subject failed to inhibit a response, the SSD was increased by a random step to increase the likelihood of success on the next stop trial. Similarly, when subjects were successful in their inhibition, the SSD was decreased to reduce the likelihood of success on the next stop trial. This procedure was employed to ensure that subjects failed to inhibit action on approximately 50% of all stop-signal trials.

On no-stop trials, the monkey was rewarded for making a saccade to the target. On stop-signal trials, the monkey was rewarded for withholding the saccade and maintaining fixation on the fixation spot. Following a correct response, an auditory tone was sounded 600ms later, and followed by a high or low fluid reward, depending on the stimulus-response mapping.

### Experimental Design & Statistical Analysis

#### Bayesian modelling of stop-signal performance

As performance on the stop-signal task can be considered as the outcome of a race between a GO and STOP process, then a stop-signal reaction time (SSRT) can be calculated (Logan and Cowan, 1984). This value can be considered as the latency of the inhibitory process that interrupts movement preparation.

Stop-signal reaction time was estimated using a Bayesian parametric approach (Matzke et al., 2013a; Matzke et al., 2013b). Compared to classical methods of calculating SSRT (i.e. integration-weighted method, Logan and Cowan (1984)), this approach allows for a distribution of SSRT to be derived by using the distribution of reaction times on no-stop trials, and by considering reaction times on non-canceled trials as a censored no-stop response time (RT) distribution. Furthermore, this model also allows for the estimation of the probability of trigger failures for a given session (Matzke et al., 2017). Individual parameters were estimated for each session. The priors were bounded uniform distributions (*μ*_*Go*_, *μ*_*Stop*_*: U* (0.001, 1000); *σ*_*Go*_, *σ*_*Stop*_*: U* (1, 500) *τ*_*Go*_, *τ*_*Stop*_*: U* (1, 500); pTF: *U* (0,1)). The posterior distributions were estimated using Metropolis-within-Gibbs sampling and we ran multiple (3) chains. We ran the model for 5000 samples with a thinning of 5.

#### EEG processing and β-burst detection

For each session, raw data was extracted from the electrode. This signal was then bandpass filtered between 15 and 29 Hz. This signal was then epoched from −1000 ms to 2500 ms relative to multiple key events in a trial, including target onset, saccade, and stop-signal presentation. β-burst detection was performed as previously described (Shin et al., 2017; Wessel, 2020). The description is adapted from therein. We then convolved the epoched signal for each trial with a complex Morlet wavelet of the form:

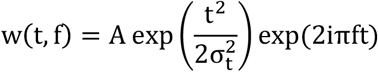

with 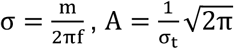, and m = 7 (cycles) for each of the 15 evenly spaced frequencies spanning the β-band (15 – 29 Hz). Time-frequency power estimates were extracted by calculating the squared magnitude of the complex wavelet-convolved data. Individual β-bursts were defined as local maxima in the trial-by-trial band time-frequency power matrix, for which the power exceeded a threshold of 6-times the median power of the entire time-frequency power matrix for the electrode. To compute the burst % across trials, we binary-coded the time of the peak β-amplitude. A β-burst density function was generated by convolving the binary-coded array of β-burst activity with a Gaussian function of the form:

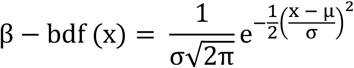

where μ = 00.0 ms, and σ = 22.5 ms. These values represent the time required for half a cycle at the median β-frequency.

#### Behavioral comparisons between monkeys

For each session, we extracted the mean response latencies on no-stop and non-canceled trials. After using the Bayesian approach described above, we extracted estimates of the stop-signal reaction time mean and standard deviation for each session. We also extracted estimates of the proportion of trigger failures for each session. Mean values between monkeys were compared using a one-way independent measure ANOVA. Greenhouse-Geiser corrections were applied when assumptions of sphericity were violated.

#### Comparing β-bursts during stopping

To examine how the β-bursts activity may vary dependent on trial type, the incidence of β-bursts observed during the stopping process were calculated for each session. The STOP process interval was defined as the time between stop-signal delay onset and stop-signal reaction time. Whilst stop trials by definition had a predefined stop-signal delay associated with them, no-stop trials were assigned a stop-signal delay value similar to that used in the most recent stop-trial. We compared this activity against a baseline period. This period was an equivalent interval of time ranging from −200 ms prior to the target onset, to −200 ms minus the mean stop-signal reaction time on the given session. For each trial type and time period (baseline & stopping), we then calculated the proportion of trials in which at least one β-burst occurred. A two-way repeated measures ANOVA was conducted, with time window and trial type as factors, and the proportion of β-bursts as the dependent variable. This approach allowed us to determine whether β-bursts were more prevalent in particular trial types, and were specific to “task-related” activity. Post-hoc tests were conducted if ANOVAs were statistically significant. To determine the time at which the incidence of β-bursts differentiated between non-canceled and canceled trials, we found the first timepoint at which the 95% confidence intervals for the β-burst density functions no longer overlapped. Greenhouse-Geiser corrections were applied when assumptions of sphericity were violated.

#### Linking incidence of β-bursts to response inhibition

To examine how neural function may reflect changes in stopping behavior, we looked at how the incidence of β-bursts varied with the probability of inhibiting a movement. This analysis was limited to stop-signal delays with 15 or more canceled trials. At each threshold, we subtracted the proportion of the β-bursts observed at a given SSD from the p (respond | stop-signal) at the same SSD. This difference between bursts and p (respond | stop-signal) was squared and values in the given session were summed, creating a sum of squared error between the two measures for each session and at each burst threshold. We then examined whether these values across sessions significantly differed from zero using a one-sample t-test at each threshold. We performed this analysis on both raw measures of β-burst proportions, and normalized measures, where incidences were relative to the maximum proportion of β-bursts observed. Findings were the same across both approaches.

#### Linking β-burst incidence to executive control

Error-related activity was examined by comparing the incidence of bursts on non-canceled trials at the middle most stop-signal delay to latency matched no-stop trials. Previous work from our lab has highlighted error-related spiking activity from this dataset became most prominent in the 100 to 300 ms period following an erroneous saccade. As such, we calculated the incidence of β-bursts that occurred during this period. Across sessions we compared the incidence of bursts observed in error trials against those observed in trials where a saccade was correctly executed using a one-way repeated measures ANOVA. Greenhouse-Geiser corrections were applied when assumptions of sphericity were violated.

#### Linking β-bursts to post-error response time adaptation

To examine how β-bursts may contribute to RT adaptations following errors or successful inhibition, we first quantified an index to capture the degree of slowing for each session. For post-error slowing, this was done by dividing the mean RT on no-stop trials following error trials by the mean RT on no-stop trials following no-stop trials for a given session. This value represents the proportional change in RT resultant from an error. We repeated this approach for post-stopping slowing, instead using the mean RT in no-stop trials following canceled trials as the numerator in this ratio.

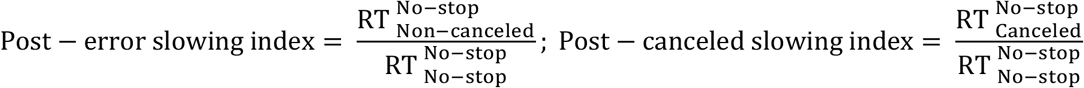

For each monkey, we examined how the incidence of β-bursts observed in the error monitoring period of a given session varied with the given sessions post-error index. To determine the association between post-error β-burst activity and post-error slowing, we fit a generalized linear model. From this we extracted R^2^ values and determined if the observed slope was significant.

We compared the effects of previous trial outcome (trial n-1) on β-burst activity observed in the baseline of the following trial (trial n). We identified no-stop trials which immediately followed canceled, non-canceled, and no-stop trials and calculated the proportion of these trials in which β-bursts occurred in the −400 to −200 ms period prior to the target appearing. We then compared if the proportion of β-bursts during this baseline period differed between different trial types using a one-way repeated measures ANOVA.

## RESULTS

We acquired 33,816 trials across 29 sessions from two macaques (Eu: 11,583; X: 22,233) performing the saccade stop-signal (countermanding) task (**Fig. 1A**). Both monkeys exhibited typical sensitivity to the stop-signal. Summary measures of performance are in **Table 1**. First, response latencies on non-canceled (error) trials were faster than those on no-stop trials (**Fig. 1B**, **left**). Secondly, the probability of failing to cancel and executing an erroneous saccade was greater at longer stop-signal delays (**Fig. 1B**, **right**). These two observations validated the assumptions of the independent race model (Logan and Cowan, 1984), allowing us to estimate the stop-signal reaction time (SSRT), the time needed to cancel to partially prepared saccade.

**Fig. 1.**
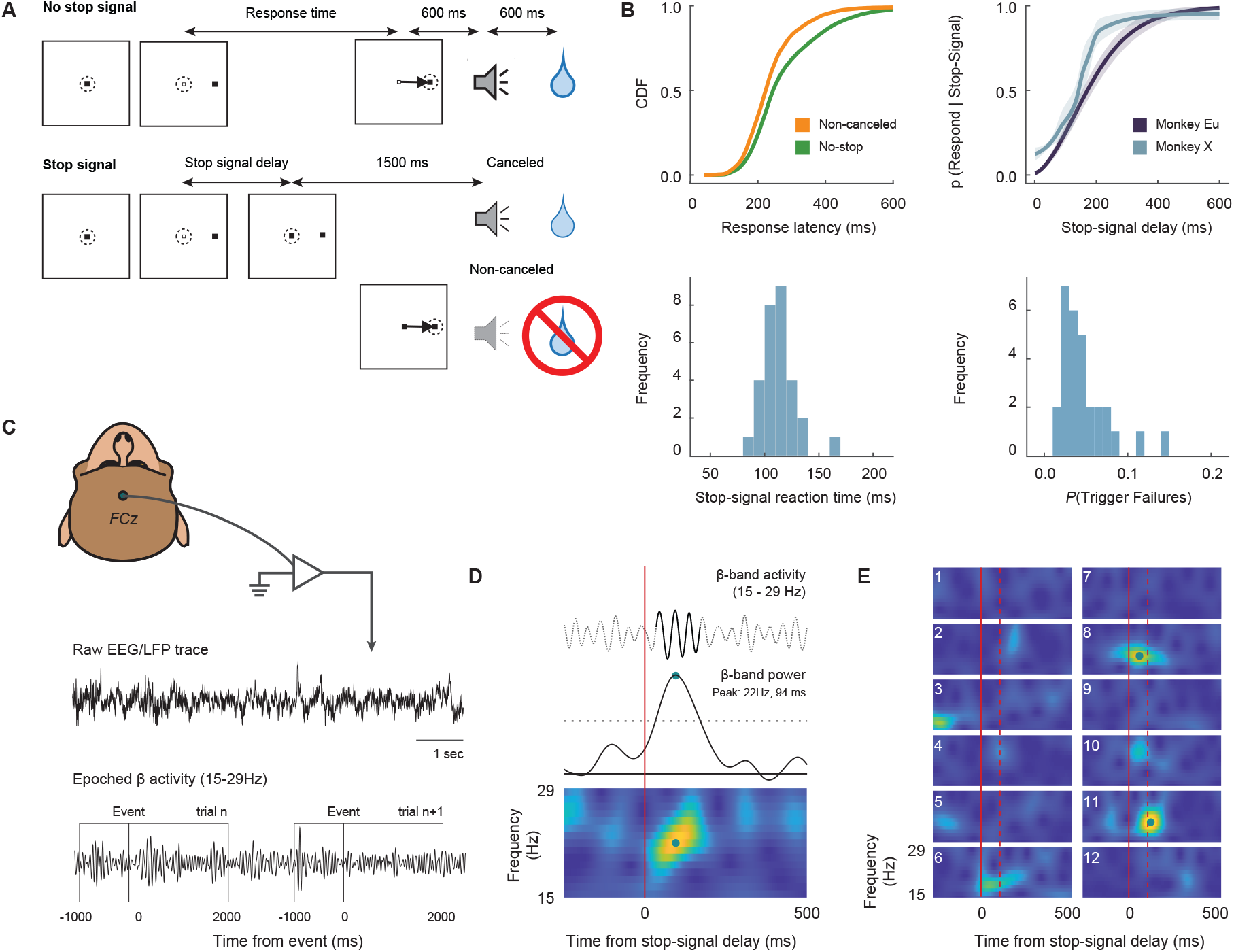
Experimental procedures. **A**. Saccade-countermanding task. Monkeys initiated trials by fixating on a central point. After a variable time, the center of the fixation point was extinguished. A peripheral target was presented simultaneously at one of two possible locations. On no-stop-signal trials monkeys were required to shift gaze to the target, whereupon after 600 ± 0 ms a high-pitched auditory feedback tone was delivered, and 600 ms later fluid reward was provided. On stop-signal trials (~40% of trials), after the target appeared the center of the fixation point was re-illuminated after a variable stop-signal delay, which instructed the monkey to cancel the saccade in which case the same high-pitched tone was presented after a 1,500 ± 0 ms hold time followed, after 600 ± 0 ms by fluid reward. Stop-signal delay was adjusted such that monkeys successfully canceled the saccade in ~50% of trials. In the remaining trials, monkeys made non-canceled errors which were followed after 600 ± 0 ms by a low-pitched tone, and no reward was delivered. Monkeys could not initiate trials earlier after errors. **B**. Countermanding behavior. *Top left*: cumulative distribution function of response latencies on no-stop (green) and non-canceled (yellow) trials. Response latencies on non-canceled trials were faster than those on no-stop trials. *Top right*: inhibition function plotting the probability of responding across stop-signal delays. Weibull functions were fitted to data from each session. The mean of these Weibull functions across sessions and the corresponding 95% CI is plotted for each monkey (monkey Eu: purple; X: blue). *Bottom left*: distribution of mean stop-signal reaction times across sessions. *Bottom right*: distribution of the proportion of trigger failures across sessions. **C**. LFP processing. Electroencephalogram (EEG) was recorded with leads placed on the cranial surface over the medial frontal cortex at the location analogous to FCz in humans. For each session, raw data was extracted. After bandpass filtering between 15 and 29 Hz, this signal was epoched from −1000 ms to 2500 ms relative to target presentation, saccade initiation, and stop-signal presentation. **D**. β-burst processing. The epoched signal for each trial was convolved with a complex Morlet wavelet. Time-frequency power estimates were extracted by calculating the squared magnitude of the complex wavelet-convolved data. Individual β-bursts were defined as local maxima in the trial-by-trial band time-frequency power matrix, for which the power exceeded a threshold of 6-times the median power of the entire time-frequency power matrix for the electrode. An example burst is shown in the time-frequency plot at the bottom. **E**. Examples of β-band time-frequency in 12 randomly selected trials. These plots are indistinguishable from counterparts derived from human data.

**Table 1:** Stop-signal task performance (mean ± SEM) for both monkeys across all sessions.

|  | **Monkey Eu**<br>**(n = 12 sessions)** | **Monkey X**<br>**(n = 17 sessions)** |
| --- | --- | --- |
| **No-stop RT (ms)** | 313.4 ± 1.6 | 263.0 ± 1.0 |
| **Non-canceled RT (ms)** | 259.4 ± 2.0 | 229.6 ± 1.0 |
| **SSRT<sub>mean</sub> (ms)** | 112.4 ± 6.4 | 114.3 ± 1.8 |
| **SSRT<sub>std</sub> (ms)** | 30.5 ± 1.8 | 30.9 ± 1.0 |
| **P (Trigger Failures)** | 0.069 ± 0.010 | 0.032 ± 0.003 |

Previous studies used SSRT for distinguishing whether neural signals can contribute directly to reactive control, so estimates of this duration must be accurate and precise. We calculated SSRT using a Bayesian parametric approach (Matzke et al., 2013a; Matzke et al., 2013b), which offers estimates of the SSRT for each session. The monkeys had indistinguishable mean SSRT (one-way independent measure ANOVA: F (1, 27) = 0.108, p = 0.745, BF_10_ = 0.367) and variance of SSRT (one-way independent measures ANOVA: F (1, 27) = 0.819, p = 0.819, BF_10_ = 0.360). This approach also quantified trigger failures when stopping was unsuccessful because the STOP process was not initialized. Trigger failures were significantly more common for monkey Eu relative to monkey X (one-way independent measures ANOVA: F (1, 27) = 18.458, p < 0.001, BF_10_ = 114.778).

### β-bursts and response inhibition

The monkeys’ electroencephalogram (EEG) was recorded with leads placed on the cranial surface over the medial frontal cortex at location analogous to FCz in humans (**Fig. 1C**). At the individual trial level, β-band activity was characterized by obvious, burst-like events, rather than by steady changes in modulations (**Fig. 1D**). As observed in human studies (Jana et al., 2020; Wessel, 2020), the overall prevalence of these bursts was low during both baseline (−400 to −200 ms pre-target, ~12.6 ± 3.8% across all sessions) and task-relevant (0 to 200 ms post-target, ~16.1 ± 5.2% across all sessions) periods.

Following previous studies (Jana et al., 2020; Wessel, 2020), to examine the relationship between β-burst activity and stopping behavior in the countermanding task we first compared the prevalence of β-bursts across trial types (two-way repeated measures ANOVA with time window and trial type as factors, Greenhouse-Geisser corrected: F (1.72, 48.31) = 9.816, p < 0.001, BF_10_ = 53.763, **Fig. 2A**, **Table 2**). We found no significant changes in the incidence of β-bursts in a baseline period and during the stop process on non-canceled trials (Holm post-hoc test, adjusted p = 0.300) or during an equivalent period of time when stopping would have occurred on no-stop trials (Holm post-hoc test, adjusted p > 0.999). However, compared to a baseline period, β-bursts were significantly more common during the STOP process when a movement was successfully canceled (Holm post-hoc test, adjusted p = 0.019). Furthermore, β-bursts were significantly more common during the STOP process on canceled compared to non-canceled trials (post-hoc test, adjusted p < 0.001) but not compared to an equivalent period of time on no-stop trials (post-hoc test, adjusted p = 0.057). This pattern of β-burst incidence replicates previous reports from human participants (Jana et al., 2020; Wessel, 2020).

**Fig. 2.**
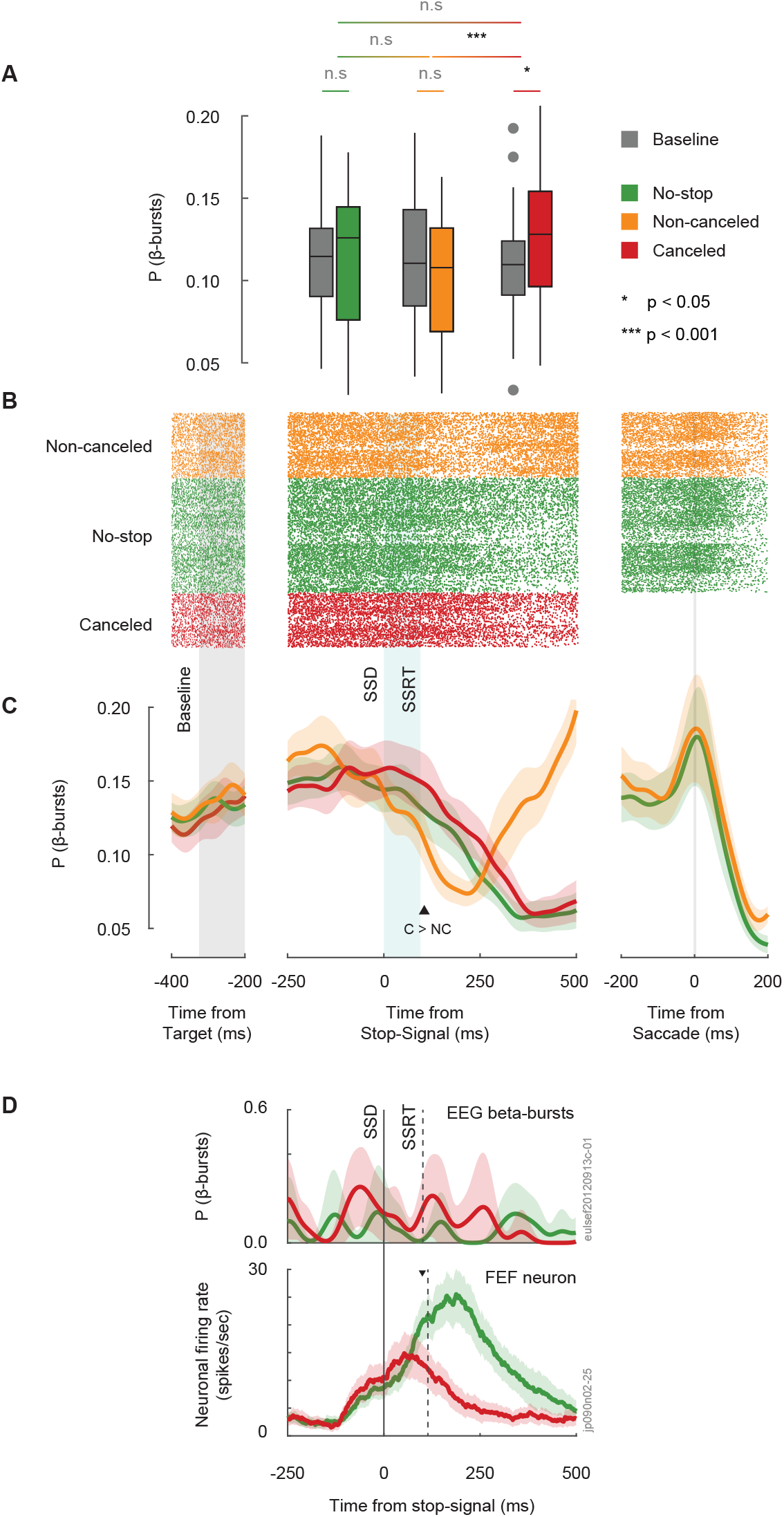
β-bursts during stopping. **A**. Boxplots showing the incidence of β-bursts observed during the STOP process interval (from scheduled SSD to SSRT) during no-stop (green), non-canceled (orange), canceled (red), and during an equivalent period of time before target presentation (grey). β-bursts are observed in ~10-15% of trials and are slightly but significantly more commonly observed when saccades are inhibited. **B**. Raster plot of β-bursts aligned on a pre-target baseline interval (left), stop signal (middle), and saccade initiation (right) across all sessions. Each tick-mark shows the time of peak β-amplitudes satisfying inclusion criteria on each trial. Raters are shown for non-canceled, no-stop, and canceled trials. The rough equivalence of β-burst frequency across types of trials is evident, as is the elevation of β-burst rate at the end of non-canceled error trials. **C**. β-burst density function derived from raster plots. β-burst peak times were convolved with a Gaussian function. During the stopping period, β-bursts were slightly but significantly more common on canceled trials (red line) than on no-stop (green) or non-canceled trials (yellow). **D**. Comparing time-course of β-burst (top) and single neuron discharges (bottom) on canceled and latency-matched no stop signal trials for a single session. At no time did the incidence of β-bursts on single session differentiate between movement initiation and inhibition. In contrast, as demonstrated previously, the discharge rate of an example FEF movement neuron sampled in one session shows a clear separation between trial types occurring before the STOP process concludes.

**Table 2:** Percentage of trials (mean ± SEM) with β-bursts during a baseline and stopping period, for all trial types.

|  | **No-stop** | **Non-canceled** | **Canceled** |
| --- | --- | --- | --- |
| **Baseline** | 11.6 ± 0.6% | 11.7 ± 0.7% | 11.2 ± 0.7% |
| **Stopping Period** | 11.8 ± 0.8% | 10.5 ± 0.7% | 13.1 ± 0.8 % |

Neurophysiological investigations quantify neural signals on a finer time scale using spike density functions (Hanes et al., 1998). To examine how changes in β-bursts occur over time, we derived β-burst density functions. First, we binary coded the time of the peak β-amplitude (**Fig. 2B**). A β-burst density function was determined by convolving this discretized array with a gaussian function over time since the stop-signal (**Fig. 2C**). These plots reveal more information about the dynamics of β-burst production through trial time and across trial types. β-burst frequency increases through the trial during saccade preparation. On no stop trials and non-canceled trials, β-burst frequency decreases following saccade initiation. The decrease in non-canceled trials begins earlier because the latency of non-canceled saccades is systematically less than that of no-stop trials. However, after noncancelled errors the incidence of β-bursts observed increases markedly. This will be characterized further below.

The incidence of β-bursts differentiated between correctly inhibited or incorrectly executed stop trials on average across sessions 132 ms after a stop-signal appeared. However, unlike the discharge rates of neurons causally involved in movement initiation and inhibition (Middlebrooks et al., 2020), β-burst density functions were so small and noisy that no time distinguished canceled from no stop trials in individual sessions **(Fig. 2D**). Notably, across sessions, β-burst incidence decreases after SSRT in canceled trials. If β-bursts are supposed to enforce response inhibition, this decay is curious. Because monkeys must sustain fixation for 1500 ms, response inhibition is sustained long after β-bursts cease.

The response inhibition function plots the fraction of non-canceled trials in which saccades are produced as a function of stop signal delay (**Fig. 1B**). The fraction of non-canceled trials is an increasing function of stop signal delay, because movements become less likely to be canceled as movement preparation progresses. Using single neuron discharge rates, a neurometric function plots the probability of modulating within SSRT as a function of stop-signal delay. The neurometric function derived from the single neuron discharges of movement-related neurons parallels the inhibition function (Brown et al., 2008). We determined whether a neurometric function derived from β-bursts parallels the inhibition function. For each session, we measured the number of β-bursts observed on canceled trials with early, intermediate, and late stop-signal delay, 50 ms before SSRT, using the conventional threshold of 6x median amplitude plus lower (2x median) and higher (10x median) thresholds. This β-burst neurometric function was compared to the probability of responding at each stop-signal delay (**Fig. 3A**) quantitatively through the sum of their squared differences at each stop signal delay. Summed squared differences close to zero indicate similarity of the two measures. Across sessions, the distribution of summed squared differences using the 6x median threshold was significantly different from zero (one-sample t-test: t (28) = 20.004, p < 0.001, BF_10_ = 1.140e+15, **Fig. 3B**). This conclusion did not depend on β-burst measurement threshold, for even with the lowest threshold where β-bursts were observed in only 40% of canceled trials. No β-burst measurement threshold produced a neurometric function similar to the inhibition function (one-sample t-test at each threshold, p < 10^−10^ for all thresholds after corrections for multiple comparisons, **Fig. 3B**).

**Fig. 3.**
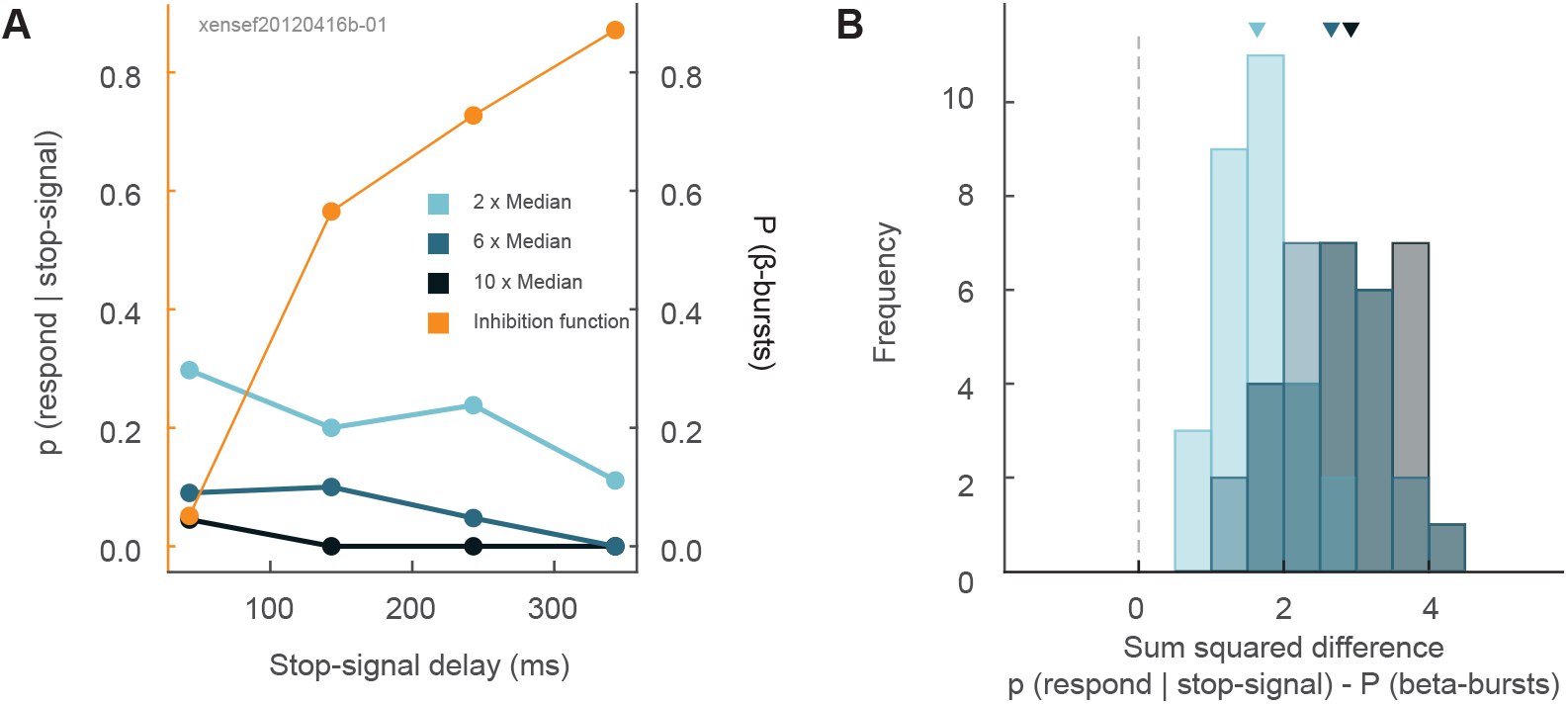
No relationship between β-bursts and response inhibition. **A**. Primary ordinate, average probability for a representative session of responding on stop signal trials as a function of stop signal delay (orange line). Secondary ordinate, average probability of β-bursts determined with amplitudes exceeding the median threshold by 2 (light blue), 6 (blue), and 10 (dark blue) in that session. The neurometric function derived from the probability of β-bursts does not correspond to the probability of responding as a function of stop signal delay. **B**. Histograms of the sum of squared differences between probabilities of canceling and of β-bursts across sessions determined with amplitudes exceeding the median threshold by 2 (light blue), 6 (blue), and 10 (dark blue). From generous to severe measurement thresholds, the incidence of β-bursts did not account for probability of cancellation with stop signal delay.

### β-bursts and executive control

The stop-signal task is useful for exploring performance monitoring because, by design, errors occur in 50% of stop-signal trials (here, 40% of all trials). Using this task, previous work has demonstrated neural activity in SEF that occurs following errors, the magnitude of which is predictive of changes in response latencies in the following trial (Stuphorn et al., 2010; Sajad et al., 2019). Hence, we compared the incidence of β-bursts 100-300 ms after error and correct saccades, when spiking activity related to errors is maximal. β-bursts were significantly more prevalent on error trials (11.1 ± 0.7%) compared to correct trials (6.7 ± 0.5%) during this period (one-way repeated measures ANOVA: F (1, 28) = 55.103, p < 10^−3^, BF_10_ = 9068.665, **Fig. 4A**).

**Fig. 4.**
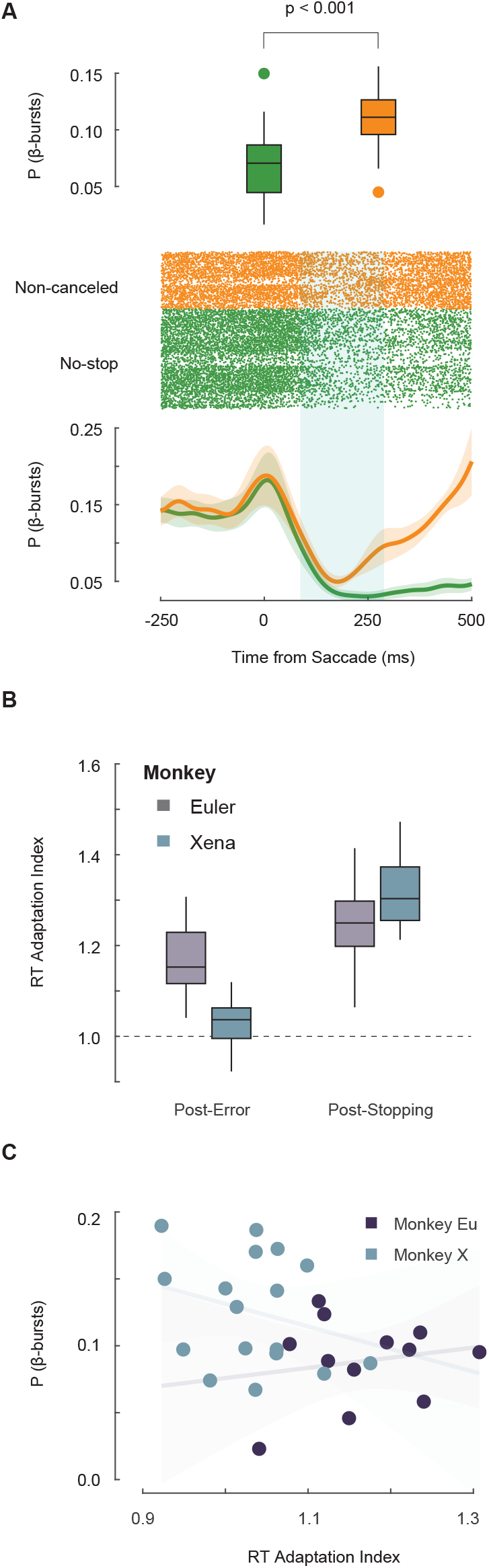
Relationship between β-bursts and performance monitoring. **A**. β-bursts during error and correct responses. *Top*: Boxplot showing significantly greater incidence of β-bursts observed 100 to 300 ms following non-canceled (orange) compared to correct saccades (green). *Middle*: Raster plot aligned on saccade. Each tick-mark shows the time of one β-burst on a non-canceled (yellow) and no-stop (green) trials. *Bottom*: β-burst density function. Following a saccade, the incidence of bursts on both error and correct trials decreases. This is followed by a pronounced increase in β-burst frequency ~150 ms after noncancelled error saccades. **B**. Boxplots of RT adaptation following non-canceled errors and successful cancellations across sessions for Eu (purple) and X (cyan). Values greater than one represent slowing. As observed previously, both monkeys tend to delay responses somewhat after errors and more after successful stopping. **C**. β-bursts are unrelated to RT adaptation. The incidence of β-bursts observed after errors did not vary as a function of the post-error RT adaptation index across sessions for Eu (purple) or X (cyan). Non-significant regression lines (± 95% CI) include 0 slope.

Behaviorally, RT on a trial varies according to the outcome of the previous trial (Emeric et al., 2007). Both monkeys produced longer RT in no-stop trials following erroneous non-canceled trials (one-way repeated measures ANOVA with previous trial type as factor, Greenhouse-Geisser corrected, F (1.54, 43.30) = 226.341, p < 0.001, BF_10_ = 6.785e+52, Holm post-hoc comparison between no-stop and non-canceled: p < 0.001, BF_10_ = 2.273e+7). To examine if post-saccade β-bursts influence post-error slowing, we calculated a post-error slowing index for each session by dividing the mean RT on no-stop trials following non-canceled trials, by the mean RT on no-stop trials following another no-stop trial (**Fig. 4B**). This value measures the relative change in RT on no-stop trials dependent on the previous trial type. We found that the incidence of β-bursts after errors bore no relation to RT on the following trial for either monkey (monkey Eu: R^2^ = 0.0333, p = 0.571, BF_10_ = 0.521; monkey X: R^2^ = 0.0732, p = 0.294, BF_10_ = 0.624, **Fig. 4C**).

## DISCUSSION

We found that the prevalence and timing of the β-bursts do not account for the likelihood of canceling a planned response. Mirroring findings with human participants, we found macaque monkeys exhibit small but consistent pulses of β-activity recorded in EEG over medial-frontal cortex during the inhibition of prepared movements (Jana et al., 2020; Wessel, 2020). Similar to these previous findings, we also observed that β-bursts occurred infrequently (~15% of trials) and were also common in trials in which a movement was generated. Furthermore, β-bursts were equally common across stop-signal delays and did not reflect changes in stopping behavior, observed in the inhibition function. Indeed, if β-bursts were to be involved in response inhibition, they should be more prevalent at earlier SSD’s where inhibition is more successful. This relationship between neurometric and psychometric measures of response inhibition has been observed in the discharges of movement-related neurons in frontal eye fields (Brown et al., 2008), but was not observed in this study. Collectively and unfortunately, these observations raise doubts about the proposal that β-bursts are a causal mechanism of response inhibition, and limit future applications in devices such as brain-machine interfaces.

Previous findings of β-bursts over frontal cortex have been interpreted as part of a larger framework proposing that the inferior frontal gyrus of the right hemisphere (rIFG) and the pre-supplementary motor area (pre-SMA) contribute to reactive control through the hyper-direct pathway (Aron and Poldrack, 2006; Aron et al., 2007). Whilst the rIFG has been implicated in the attentional capture of the stop-signal & initiating the STOP process (Swann et al., 2012; Jana et al., 2020), the role of the pre-SMA is less clear. Human fMRI studies show greater activity in pre-SMA on stop trials with manual responses (Aron and Poldrack, 2006; Aron et al., 2007; Rae et al., 2015) and in supplementary eye field with saccades (Thakkar et al., 2014). Lesion studies also associate medial frontal areas with impaired stopping of limbs (Floden and Stuss, 2006; Nachev et al., 2007; Sumner et al., 2007) and eyes (Husain et al., 2003). This is mirrored in human electrophysiological evidence reporting stronger signals over pre-SMA during canceled trials (Swann et al., 2012). However, our observation that β-bursts are slightly common during response inhibition is unexpected in macaques based on previous neurophysiological results. In single-unit recordings, the modulation of neurons in MFC occurs too late to contribute to the reactive control of movement and instead contributes to performance monitoring and the exertion of proactive control (Emeric et al., 2010; Stuphorn et al., 2010). Such discrepancies have sparked debate about whether macaques are a useful model of response inhibition and performance monitoring in humans (Cole et al., 2009; Schall and Emeric, 2010). However, this question is confounded by different response modalities, tasks, and measurement scales. We addressed the problem of measurement scale directly in this study.

Whilst we found β-bursts were more common during response inhibition, we also found they were more common following errors, suggesting a contribution to performance monitoring and executive control. This observation is consistent with patterns of spiking activity described in medial frontal cortex (Stuphorn et al., 2000; Sajad et al., 2019) and complementing previous observations that error, conflict, and reward monitoring are typically associated with theta (4-8 Hz) band modulation (Luu et al., 2004; Trujillo and Allen, 2007; Cohen et al., 2008; Cavanagh et al., 2009; Nigbur et al., 2011; Amarante et al., 2017). Over medial frontal cortex, β-power is elevated when cognitive control was required and following negative feedback (Stoll et al., 2016). Specifically, in a change-stop-signal task, previous work has highlighted greater β-activity on canceled compared to non-canceled trials in pre-SMA, following the completion of the STOP process (Jha et al., 2015). We also found that the incidence of β-bursts after errors did not predict post-error adjustments in RT. This contrasts with a recent observation using MEG focused on pre-SMA (Jha et al., 2015), but is in line with the inconsistent and variable relationship between the error-related negativity and performance adjustments (Gehring et al., 1993; Gehring and Fencsik, 2001; Rodriguez-Fornells et al., 2002; Hajcak et al., 2003; Kerns et al., 2004; Holroyd et al., 2005; Ladouceur et al., 2007; West and Travers, 2008; Nunez Castellar et al., 2010; Godlove et al., 2011b; Reinhart et al., 2012; Fu et al., 2019).

To conclude, by replicating measurements of β-bursts in EEG of macaque monkeys, we establish an animal model of this phenomenon. First, we demonstrated β-bursts were more prevalent when movements were successfully inhibited. Second, we demonstrated a greater incidence of β-bursts over the medial frontal cortex when a movement was erroneously executed. However, in neither context were β-bursts frequent enough to account for behavior. Given the pulses of β-bursts at different stages during the task, it is uncertain whether they index different mechanisms or are produced by one mechanism at different times. This uncertainty and the apparent absence of causal efficacy have been explained by appeals to the poor signal to noise ratio of noninvasive EEG recordings. Establishing an animal model affords systematic investigation of the neural mechanisms of β-burst generation in the cerebral cortex during countermanding or other tasks. By sampling β-bursts across the layers of the cortex, we can provide insights into the mechanisms of their generation and possibly elucidate the role of β-bursts during response inhibition and executive control.

## ACKNOWLEDGEMENTS

We thank B. Williams, R. Williams, M. Maddox, M.S. Schall, B. Haniff, S. Motorny, D. Richardson, L. Toy, and M.R. Feurtado, for technical support. We also thank K.A. Lowe, Dr U. Rutishauser, Dr. A. Sajad, J.A. Westerberg, and Dr. T. Womelsdorf, for useful conversations regarding the work.

